# Pregranulosa cells engage a distinct transcriptional programme prior to cell-cycle dependent primordial follicle activation

**DOI:** 10.1101/2022.10.24.513438

**Authors:** Güneş Taylor, Emily R Frost, Brendan N Crow, Arthur Radley, Stefan Boeing, Christophe Galichet, Barbora Bucinskaite, Mark A Baker, Jessie M Sutherland, Robin Lovell-Badge

**Affiliations:** Laboratory of Stem Cell Biology and Developmental Genetics, The Francis Crick Institute, London, United Kingdom, NW1 1AT; School of Biomedical Science & Pharmacy, University of Newcastle, Ring Road, Callaghan, New South Wales, Australia, 2308; Hunter Medical Research Institute, New Lambton Heights, New South Wales, Australia, 2305; Laboratory of Developmental Dynamics, The Francis Crick Institute, London, United Kingdom, NW1 1AT; Bioinformatics and Biostatistics Facility, The Francis Crick Institute, London, United Kingdom, NW1 1AT; Scientific Computing – Digital Development Team, The Francis Crick Institute, London, United Kingdom, NW1 1AT; Centre of Reproductive Health, Institute of Regeneration and Repair, University of Edinburgh, Scotland, EH16 4UU; Neurobiological Research Facility, Sainsbury Wellcome Centre, London, United Kingdom, W1T 4JG

**Author notes:** Corresponding authors: *Dr Güneş Taylor*, Centre of Reproductive Health, University of Edinburgh, Scotland, EH16 4UU, *Prof Robin Lovell-Badge*, Stem Cell Biology and Developmental Genetics Lab, The Francis Crick Institute, London, United Kingdom, NW1 1AT, *Dr Jessie Sutherland*, School of Biomedical Science & Pharmacy, University of Newcastle, Ring Road, Callaghan, New South Wales, Australia, 2308. These authors contributed equally and should be regarded as co-first authors (Güneş Taylor, Emily R Frost). These authors contributed equally and should be regarded as co-last authors (Jessie M Sutherland, Robin Lovell-Badge).

**Keywords:** ovary, oocyte, infertility, granulosa cells, cell cycle, cdkn1b, p27, TNNI3, primordial follicle activation, entropy sorting

## Abstract

Primordial follicles are quiescent ovarian structures comprised of a single oocyte surrounded by a layer of somatic supporting pregranulosa cells. Primordial follicle activation is the first step towards oocyte maturation and, ultimately, ovulation. As the number of quiescent primordial follicles is finite, their rate of activation is a critical parameter of the duration of the female reproductive lifespan. Activation status is established by the presence of cuboidal and proliferative granulosa cells in primary follicles, rather than squamous and quiescent pregranulosa cells in primordial follicles. Here, using a continuous Entropy Sort Feature Weighting approach on single-cell RNA sequencing data, we identify a distinct transcriptomic signature of activating pregranulosa cells in neonatal wildtype mice. This signature contains several genes previously linked with mature granulosa cells as well several novel candidates: *Slc18a2, Tnni3, Fam13a* and *Myo1e*. We confirm expression of *Slc18a2* and TNNI3 in the granulosa cells of activating follicles in embryonic, neonatal and adult mouse ovaries. Perturbation of cell cycle inhibitor p27^kip1^ in *Cdkn1b*^-/-^ mice results in complete activation of all primordial follicles during this neonatal period. Contrary to previous reports on this established mouse model, we find substantial transcriptomic changes in embryonic *Cdkn1b*^-/-^ ovaries. Upon loss of cell-cycle inhibition, we find increased expression of our signature of pregranulosa cell activation, particularly that of cardiac troponin I (*Tnni3*). We conclude that pregranulosa cells engage a distinct transcriptional programme prior to cell-cycle dependent primordial follicle activation.

## Introduction

Critical parameters of the female reproductive lifespan are the finite number of quiescent primordial follicles within their ovaries and the rate of primordial follicle activation for maturation and eventual ovulation. Primordial follicles are formed during gestation and consist of an oocyte with a single layer of surrounding pregranulosa cells (1, 2). Once primordial follicles activate, the pregranulosa cells become steroidogenic granulosa cells and provide structural and metabolic support to the oocyte throughout folliculogenesis (3). Primordial follicles are initially formed and distributed throughout the ovary. However, typically during late foetal development or neonatally in mammals, the majority of follicles within the central ovarian medulla are lost subsequent to a synchronised activation process termed first wave activation (4, 5). In rodents, the sex steroids produced by supporting cells during first wave activation are hypothesised to be important for establishing aspects of puberty and the ovarian–hypothalamic–pituitary axis (4, 6, 7). Following first wave activation, the remaining primordial follicles reside in the outer ovarian cortex and are cyclically activated for eventual ovulation or atresia. The establishment of this ‘ovarian reserve’ of cortical primordial follicles, and their gradual release from it, sustains fertility over many months in mice and several decades in humans (8).

Consequently, stringent control of primordial follicle activation is required to ensure that the ovarian reserve persists for the entire reproductive lifespan. Key indicators of primordial follicle activation are changes in granulosa cell morphology (from squamous to cuboidal) and re-entry into the cell-cycle, and increased oocyte size (9-12). The morphological changes to pregranulosa cells precede that of the oocyte, suggesting that primordial follicle activation may be initiated by the pregranulosa cells (13-15). While many pathways have roles in follicle activation, including PI3K, mTOR, KITL, TGFB and Hippo signalling (9, 10, 16-19), the signal initiating primordial follicle activation remain unresolved. Primordial follicle activation is challenging to interrogate due to the asynchronous nature of activation throughout the adult ovarian cortex. Advances in single-cell RNA sequencing (scRNAseq) have accelerated our understanding of many ovarian processes in the mouse (20-22), however the difficulty of knowing which cortical follicle is initiating activation has impeded the identification of a specific gene or process involved in the earliest steps of this critical process.

Here we leveraged the abundance of activating follicles during first wave activation to investigate the transcriptomic changes of pregranulosa cells in neonatal mice. We identify a putative signature of activating pregranulosa cells by analysing scRNAseq data using continuous Entropy Sort Feature Weighting (cESFW). We confirm that two novel targets within the signature, solute carrier family 18 member 2 (*Slc18a2*) and cardiac troponin I (*Tnni3*), increase in expression postnatally within transitioning pregranulosa cells and granulosa cells. Furthermore, we find increased expression of our signature genes within the ovaries of a mutant line with precocious and complete first wave activation (*Cdkn1b*^-/-^ or *Cdkn1b* / p27^kip1^-null) (23). Contrary to previous reports, we also find that loss of *Cdkn1b* results in transcriptomic changes at E18.5, prior to the morphological changes indicative of follicle activation. We confirm *Tnni3* mRNA expression in *Cdkn1b*^-/-^ granulosa cells at E18.5. These data demonstrate that pregranulosa cell maturation into granulosa cells is a key driver of primordial follicle activation and integrates cell-cycle regulator *Cdkn1b* / p27^kip1^ within the molecular mechanism driving this important cell transition.

## Results

### Identification of putative regulators of primordial follicle activation within pregranulosa cells

To profile the transcriptomic dynamics of first wave activation in wildtype (WT) mice, scRNAseq was conducted on ovaries at embryonic day 18.5 (E18.5), post-natal day 4 (PD4), and post-natal day 7 (PD7) (Figure 1a). These timepoints captured prior to, during and after first wave activation respectively. Using our previously described protocol (24), we enriched for the somatic cell compartment in the single-cell suspension. The dataset contained 24,810 cells in total with 8,210 cells at E18.5, 7,587 cells at PD4 and 9,013 cells at PD7. Data from all three timepoints were integrated and, subsequent to default seurat highly variable gene selection, uniform manifold approximation and projection (UMAP) plots were generated with 18 distinct clusters. The six major cell types (granulosa cells Gc_1 – Gc_5), mesenchymal cells (Mc_1 – Mc_5), oocytes (Oo_1), epithelial cells (Ep_1 – Ep_2), immune cells (Im_1) and blood cells (Bl_1 – Bl_2)) in the ovary were annotated based on their expression of established cell type markers (Figure 1a and Supplemental Figure 1a) (5).

**Figure 1.**
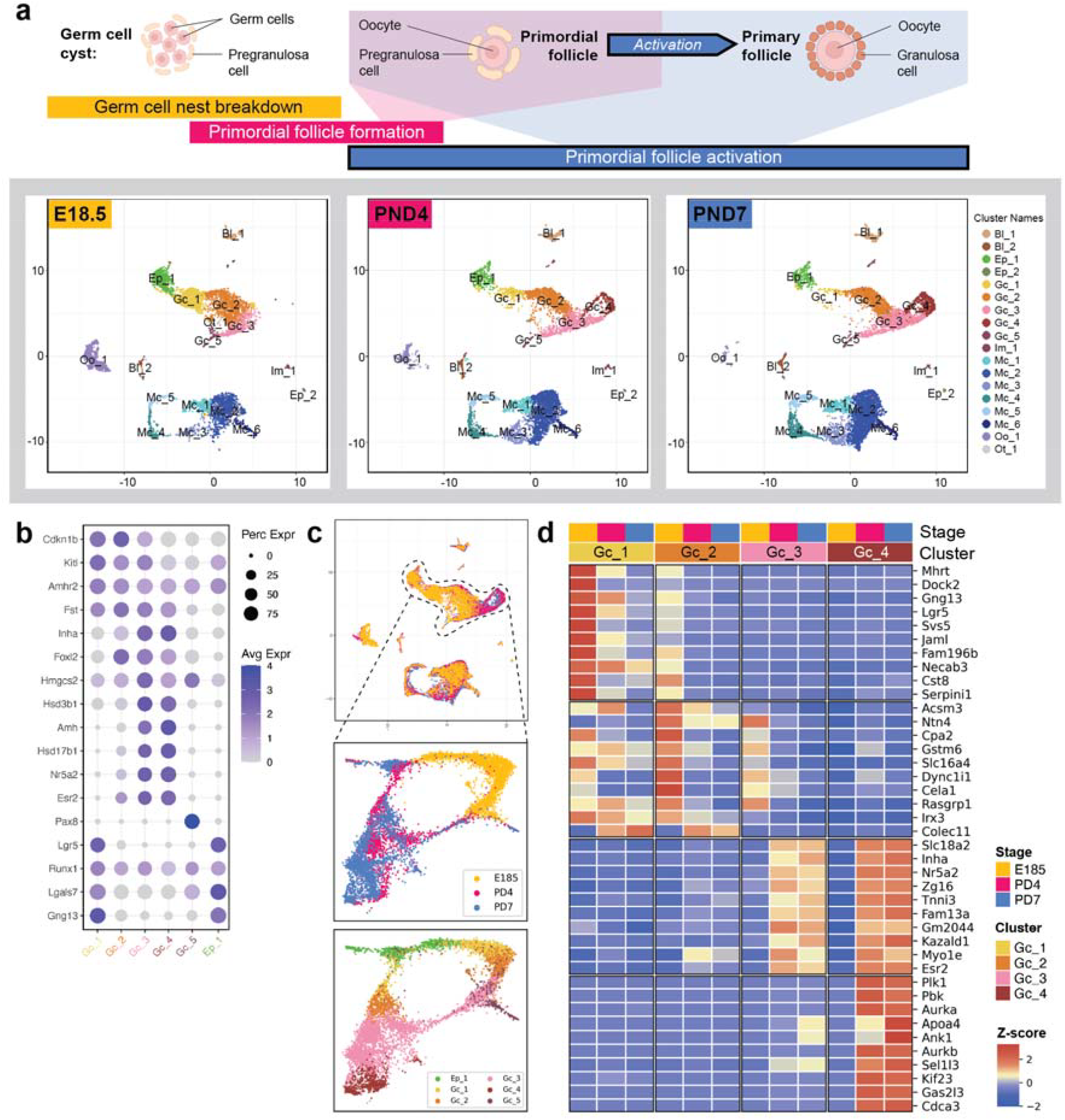
scRNAseq of mouse ovaries reveals transcriptomic differences in pregranulosa cells postnatally. **a** UMAPs from scRNAseq on whole mouse ovaries before (E18.5), during (PD4) and after (PD7) primordial follicle activation. 18 clusters representing major ovarian cell types were identified. **b** Dotplot showing known markers of pregranulosa cells are found in Gc_1, Gc_2, Gc_3 and Gc_4. **c** UMAP from cESFW workflow. Middle panel: coloured by developmental timepoints - E18.5 (yellow), PD4 (magenta) and PD7 (blue). Bottom panel: Unsupervised clusters (Supplemental Figure 1d) coloured by cell type labels from conventional scRNAseq analysis workflow (Ep_1, Gc_1, Gc_2, Gc_3, Gc_4 and Gc_5). **d** Heatmap top 10 entropy sort score genes within three pregranulosa cell clusters (Gc_1, Gc_2, Gc_3) and granulosa cells (Gc_4).

We focused on discovering differences between Gc_1 through Gc_4 as cluster Gc_5 had high expression of *Pax8* (Figure 1b), a recently reported marker of supporting-like cells from the *rete ovarii* (25). Using a detailed panel of pregranulosa and granulosa cell markers selected from the literature, we classified Gc_1 as an embryonic population of epithelial-derived pregranulosa cells due to their expression of *Lgr5, Gng13* and *Lgas7* (Figure 1b), and the decreased cell numbers in Gc_1 between E18.5 and PD7 (Figure 1a) (4, 5, 26, 27). We identified Gc_2 as a population of pregranulosa cells derived from the Nr5a1+ somatic cell lineage based on their high expression of *Cdkn1b* and *Foxl2* alongside low expression of *Hsd3b1* and *Hmgcs2* (Figure 1b) (5). Interestingly, Gc_3 had similar gene expression to Gc_2 (shown by the expression of *Foxl2* and *Hmgcs2* amongst others), but with elevated expression of well-established markers of activated granulosa cells like *Kitl* and *Fst* (Figure 1b) (10, 28). This suggested that Gc_3 is a population of pregranulosa cells that have initiated primordial follicle activation. Cluster Gc_4 expressed mature granulosa cell marker genes including *Amhr2* and *Hmgcs2* (Figure 1b), were in the G2M phase of the cell cycle (Supplemental Figure 1b) and showed high expression of proliferation genes including *Top2a* and *mKi67* (9, 29). When interrogated by timepoint and cell cycle phase, we observed that clusters Gc_1, Gc_2 and Gc_3 were present from E18.5, while Gc_4 only appeared at PD4 during first wave activation and was in the proliferative G2M / S phase of the cell cycle (Supplemental Figure 1b). Overall, the gene expression changes in mature granulosa cell and proliferation genes across Gc_1 – Gc_4 allowed for the separation of granulosa cell populations into distinct clusters. This delineation provided the opportunity to delve into the granulosa cell transcriptome across the primordial follicle transition into folliculogenesis.

To interrogate the granulosa cell populations more closely, we subsetted the Gc_1 - Gc_5 clusters Gc_1 - Gc_5, along with the Ep_1 cluster, the surface epithelium from which many supporting cells are derived (Supplemental Figure 1b and d). Overall, two distinct gene expression dynamics emerged: genes that increase in expression during activation and persist in mature granulosa cells (Gc_3 and Gc_4), and genes that decreased upon entry into the activating pregranulosa cell pool (Gc_1 and Gc_2) (Figure 1b). Differential expression analysis showed there was significant overlap in gene expression across all Gc clusters, and highlighted that distinct marker genes exist for Gc_4 but not the remaining Gc clusters (Supplemental Figure 1c, Dataset S1-4). As an alternative to highly variable gene selection analysis for feature selection, we applied a new computational approach called ‘continuous Entropy Sort Feature Weighting’ (cESFW) (30) to the Ep_1 and Gc_1 – Gc_5 scRNAseq clusters. Subsetting the scRNA-seq data to the 4376 top ranked genes identified by cESFW generated a higher resolution UMAP of granulosa cell development (Figure 1c). Unsupervised Kmeans clustering revealed 14 distinct clusters (Supplemental Figure 1d). To compare against the differential expression analysis, we manually assigned the cESFW Kmeans clusters to the cell type annotations identified using the conventional scRNAseq workflow (Figure 1c). We then used entropy sorting (ES) (30) to calculate the entropy sort score (ESS) correlation metric to get ranked gene lists for each of these clusters (Figure 1d, Dataset S5). The top ten markers of each cluster differed from those found by default differential gene expression analysis but also revealed, once upregulated Gc_3 markers are maintained in Gc_4. As such, cESFW feature selection separates the transcriptional signature of the onset of pregranulosa cell activation (e.g. *Slc18a2, Nr5a2, Fam13a* and *Tnni3*) from that of mature granulosa cells only (e.g. *Aurka, Kif23*).

### Dynamic transcriptome changes underpin the transition of pregranulosa cells to granulosa cells

To ensure that our signature genes from cESFW were specific to activating pregranulosa cells, we selected two targets for further analysis. *Tnni3* encodes for an actin-binding protein involved in the troponin complex in cardiac tissues and was recently reported in the granulosa cells of maturing follicles in humans (31, 32). Solute carrier family 18-member A2 (*Slc18a2*) is a vesicular monoamine transporter, associated with FSH levels in patients with polycystic ovary syndrome (33). We probed WT mouse ovaries at E18.5, PD4 and PD7 for *Slc18a2* RNA expression and TNNI3 protein expression (Figure 2a and Figure 2b). *Slc18a2* was not detected within ovarian sections at E18.5 (Figure 2a). However, at PD4 and PD7, *Slc18a2* foci were apparent within cuboidal granulosa cells (Figure 2a). Similarly, TNNI3 staining was low to undetectable within pregranulosa cells at E18.5, but at PD4 and PD7, TNNI3 staining was apparent within granulosa cells (Figure 2b). Additionally, we captured transitioning follicles (containing both cuboidal granulosa cells and pregranulosa cells) where *Slc18a2* and TNNI3 staining was only observed in the activated granulosa cells of a single follicle (Figure 2c and Figure 2d). Throughout the ovaries, staining for TNNI3 was seen in oocytes of growing follicles but minimal staining was seen in the oocytes of primordial follicles (Figure 2d). This was reflected in the scRNAseq data with transcription of *Tnni3* in 11.6% of cells in the oocyte cluster. We also confirmed that granulosa cell expression of TNNI3 was conserved across timepoints, detecting staining in activated cortically derived follicles in adult mouse ovaries (Supplemental Figure 2). This illustrates upregulation of TNNI3 both in medullary follicles activated during first wave activation and in the cortical follicles of the adult. Taken together, our data show that pregranulosa cells upregulate *Slc18a2* and *Tnni3* expression upon entry into follicle activation. This expression pattern is conserved across the two waves of follicle activation, in both the juvenile and adult mouse ovary for TNNI3 (Supplemental Figure 2).

**Figure 2.**
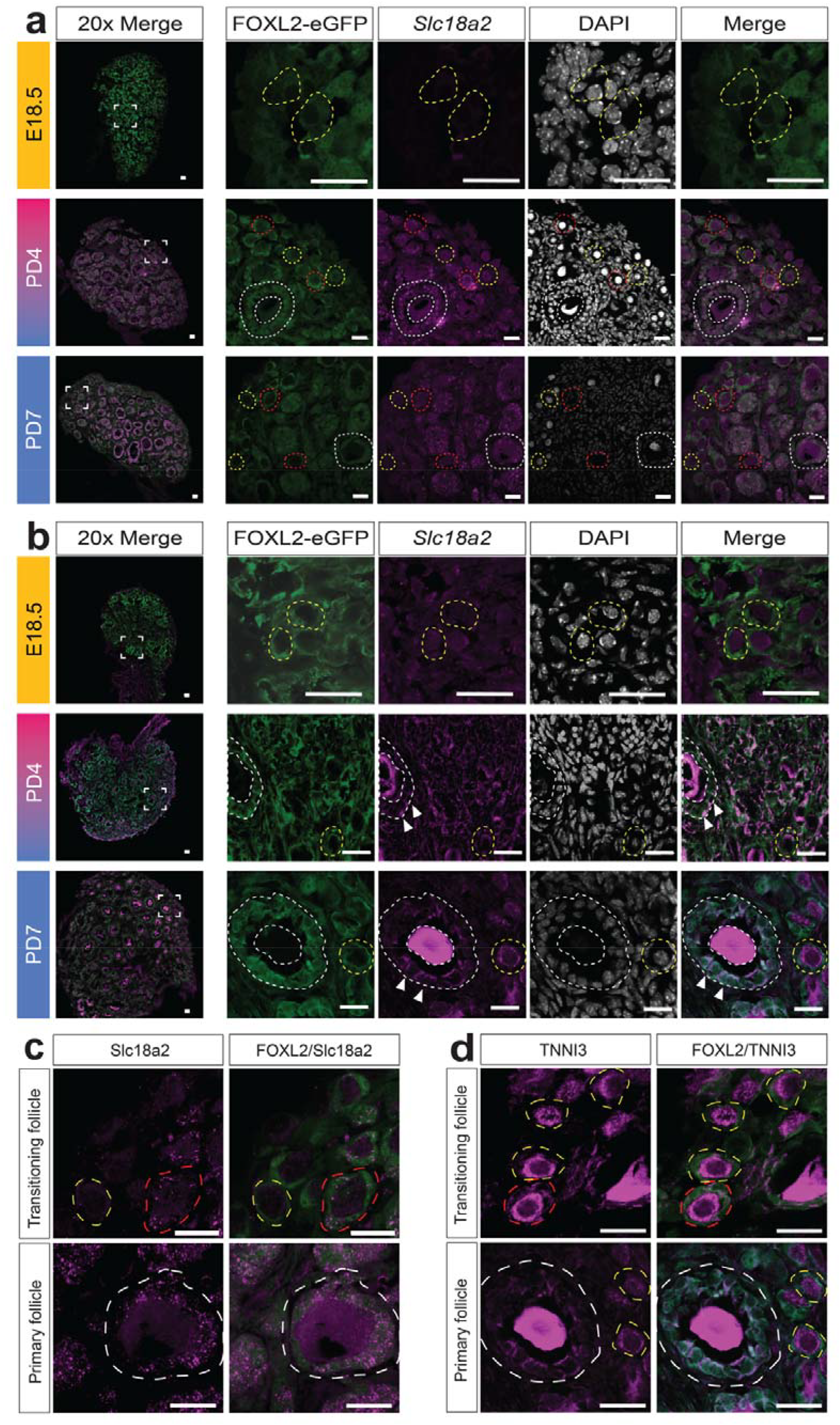
Pregranulosa cells show a distinct gene expression signature as they activate. **a** In situ hybridisation and immunofluorescence on *Foxl2*^p2A-eGFP^ ovarian cryosections at E18.5 (n=3), PD4 (n=3) and PD7 (n=3). Each section contains eGFP (green) labelling in all granulosa and pregranulosa cells, a HCR probe for *Slc18a2* (magenta) and DAPI (white). 20x magnification images shown in left column of each timepoint. E18.5 samples were further imaged at 100x magnification. PD4 and PD7 samples were additionally imaged at 63x magnification. Growing follicles are denoted by a white dashed border while primordial follicles are denoted by a dashed yellow border. Transitional follicles denoted in red dashed border. White scale bars show 20µm. **b** Immunofluorescence on *Foxl2*^p2A-eGFP^ ovarian cryosections at E18.5 (n=3), PD4 (n=3) and PD7 (n=3). Each section is immunolabeled for GFP (green) labelling all granulosa and pregranulosa cells, TNNI3 (purple) and DAPI (white). 20x magnification images shown in top row of each timepoint. E18.5 samples were further imaged at 100x magnification. PD4 and PD7 samples were additionally imaged at 63x magnification. Growing follicles are denoted by a white dashed border while primordial follicles are denoted by a dashed yellow border. White scale bars show 20µm. **c** High magnification image of an PD7 *Foxl2*^p2A-eGFP^ transitional follicle (red dashed line) displaying high punctate *Slc18a2* staining throughout the granulosa cells (indicated by white arrows). Scale bars = 20µm. **d** High magnification image of an PD7 *Foxl2*^p2A-eGFP^ transitional follicle (red dashed line) displaying high TNNI3 staining predominantly around the edge of the granulosa cells (indicated by white arrows). Scale bars = 20µm.

### Pregranulosa cells undergo transcriptomic changes precociously in *Cdkn1b*^-/-^ mice

To further explore the involvement of the genes putatively underlying pregranulosa cell activation, we queried if our activation signature genes are disrupted in an established model of precocious and aberrant follicle activation, *Cdkn1b*^-/-^ (*Cdkn1b* / p27^kip1^-null) mice. *Cdkn1b* / p27^kip1^ is a cyclin dependent kinase inhibitor and therefore blocks cell cycle progression. *Cdkn1b*^-/-^ mice show uncontrolled activation of follicles perinatally, inducing infertility prior to sexual maturation (23, 34). We confirmed that populations of germ cells and granulosa cells are present with *Cdkn1b*^-/-^ ovaries using DDX4 and FOXL2 staining, as seen previously (Supplemental Figure 3a) (23, 34). We performed bulk RNA-sequencing of control and *Cdkn1b*^-/-^ ovaries at three timepoints: E13.5, E18.5 and PD8 (Supplemental Figure 3b, Dataset S6). As ovarian supporting cells express *Cdkn1b* / p27^kip1^ prior to expression of the pregranulosa cell specification marker *Foxl2*, the E13.5 timepoint was chosen to confirm pregranulosa specification is unperturbed by the loss of *Cdkn1b* (12, 35). The E18.5 timepoint was chosen as it is prior to first wave activation in WT mice and is the first timepoint of our scRNAseq timecourse. First wave activation has concluded by PD8 and was investigated in the previous report of *Cdkn1b* / p27^kip1^-null mice and aligns with our final scRNAseq timepoint.

We found just 4 differentially expressed genes at E13.5 (p < 0.05) (*Cdkn1b, Eya4, Pitx2* and *Fcnb*), confirming pregranulosa cell specification is essentially unperturbed. As previously reported, larger numbers of primary and secondary follicles were observed in *Cdkn1b*^-/-^ ovaries at PD8 (Figure 3a) and we found 746 differentially expressed genes at PD8 in our bulk RNAseq data (p < 0.05, Dataset S6). To check transcriptomic changes prior to primordial follicle activation in the wild-type ovary, we examined ovaries from both genotypes at E18.5 and unexpectedly found 896 upregulated and 520 downregulated differentially expressed genes in *Cdkn1b*^-/-^ ovaries compared to WT controls (p < 0.05) (Figure 3b, Supplemental Figure 3c and Dataset S6). Contrary to previous reports, this suggests that absence of *Cdkn1b* causes precocious first wave activation around the time of germ cell nest breakdown and primordial follicle formation.

**Figure 3.**
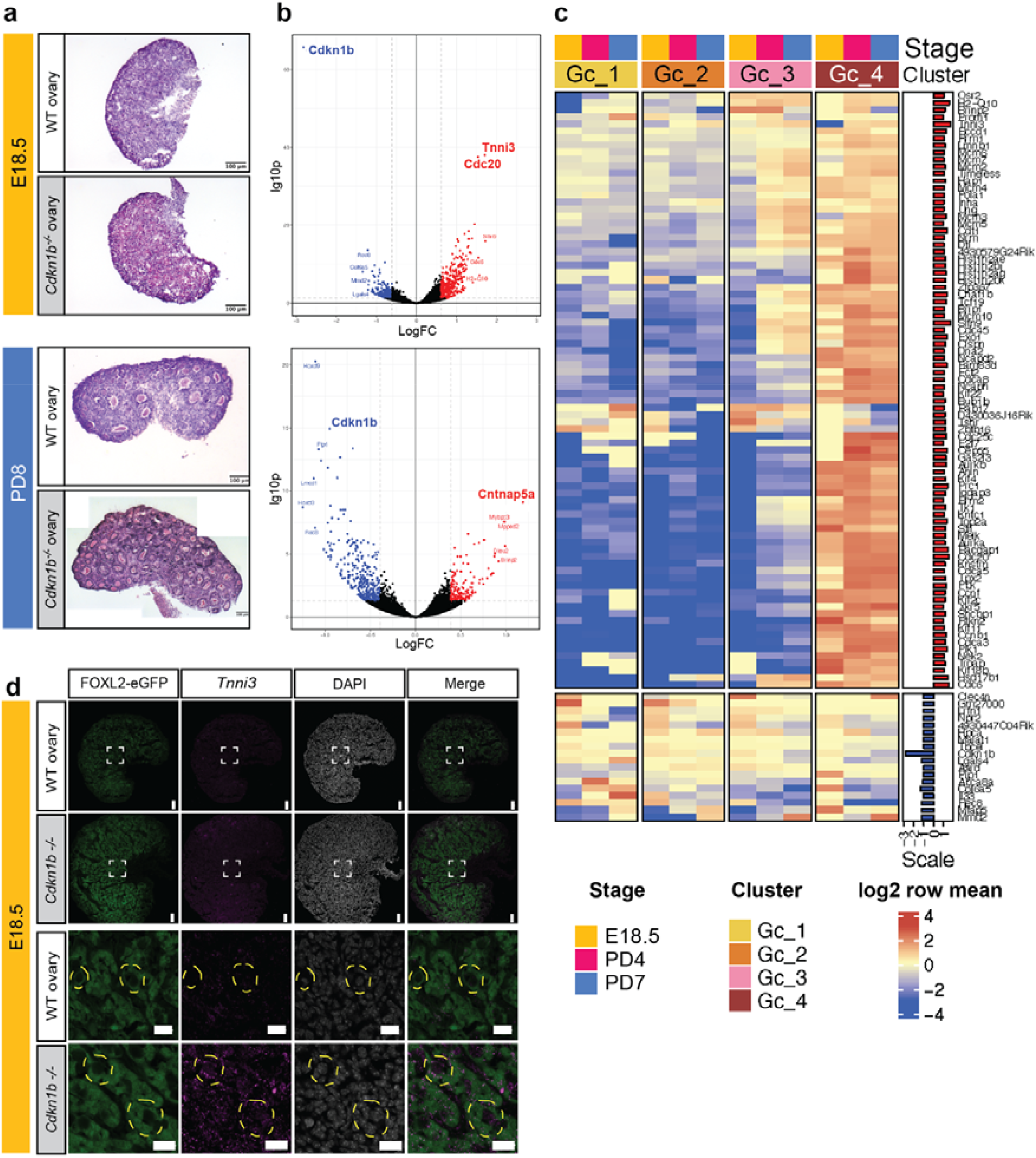
Disruption of p27^Kip1^ highlights functional aspects of activating pregranulosa cell gene expression signature. **a** Haematoxylin and Eosin staining of E18.5 and PD8 ovaries in WT and *Cdkn1b*^-/-^ mice. Imaged at 10x magnification. **b** Volcano plots showing differential expression of gene transcripts between E18.5 WT and *Cdkn1b*^-/-^ ovarian cells (upregulated genes in red, downregulated in blue, n = 4 replicates per group). Black dots indicate genes below the p ≤ 0.05 cutoff. **c** Heatmap showing differentially expressed genes in *Cdkn1b*^-/-^ ovaries (p ≤ 0.05) overlaid with granulosa cell subclustering analysis from the scRNAseq experiment (upregulated genes in red, downregulated genes in blue). **d** In situ HCR on *FoxL2*^p2A-eGFP^ (WT) and *FoxL2*^p2A-eGFP^; *Cdkn1b*^-/-^ ovarian cryosections at E18.5 (n=3). Each section is imaged for eGFP (green) labelling all granulosa and pregranulosa cells, *Tnni3* (magenta) and DAPI (white). 20x magnification images shown in top panel and 63x magnification images shown in bottom panel for both genotypes. Primordial follicles are denoted by a dashed yellow border. White scale bars show 20µm.

Since in the ovary *Cdkn1b* / p27^kip1^ is confined to pregranulosa cells at this developmental stage, we hypothesised that the specific maturation of pregranulosa cells to granulosa cells was driving this precocious primordial follicle activation. Therefore, we compared the 1416 differentially expressed genes from the bulk RNA-sequencing of E18.5 *Cdkn1b*^-/-^ ovaries with the genes expressed in clusters Gc_1 to Gc_4 from our scRNAseq timecourse (Figure 3c). This revealed that most of the differentially expressed genes in *Cdkn1b*^-/-^ ovaries at E18.5 were expressed in the activating pregranulosa cell cluster Gc_3 and granulosa cell cluster Gc_4 in WT ovaries. Importantly, the most statistically significantly upregulated differentially expressed gene at E18.5 in *Cdkn1b*^-/-^ ovaries was *Tnni3* (1.70logFC, padj=1.06E-38), a gene also identified by the ESS as one of the top ten markers of Gc_3 activating pregranulosa cells (Figure 1d). *Tnni3* expression is restricted to Gc_3 and Gc_4 cell clusters, contrary to the high *Cdkn1b* expression in clusters Gc_1 and Gc_2 (Figure 2b). *In situ* hybridisation confirmed the localisation of punctate *Tnni3* to pre-granulosa cells in both WT and *Cdkn1b*^-/-^ ovaries, with significantly higher puncta intensity (2.1x fold change) in *Cdkn1b*^-/-^ ovaries (Figure 3d). Together, this suggested engaging the cell-cycle via reduction of *Cdkn1b* expression and increased *Tnni3* expression may drive maturation of pregranulosa cells into granulosa cells in the mouse.

## Discussion

Primordial follicle activation is critical to the establishment and duration of mammalian female fertility. Understanding the molecular mechanisms underpinning activation holds the potential to improve reproductive longevity, assist in the diagnosis of infertility and develop targeted treatments for patients with ovarian dysfunction. To interrogate this mechanism, we generated a scRNAseq dataset of first-wave primordial follicle activation. We focused on four distinct clusters of granulosa cells that undergo morphological changes associated with initiation of primordial follicle activation. One cluster, Gc_3, represented a distinct population of activating pregranulosa cells, which was interrogated through entropy sorting (30). This analysis revealed a transcriptomic signature of activating pregranulosa cells, where several strong candidate genes for the initiation of follicle activation were identified, including *Slc18a2* and *Tnni3*. We confirmed that our signature contained genes altered during follicle activation using the *Cdkn1b*^-/-^ mouse line, a model for accelerated primordial follicle activation. Signature Gc_3 genes were dysregulated in *Cdkn1b*^-/-^ ovaries, including *Tnni3* which showed precocious upregulation in *Cdkn1b*^-/-^ pre-granulosa cells. These data implicated the cardiac troponin gene, *Tnni3*, as a cell-cycle regulated marker of primordial follicle activation within pregranulosa cells.

The ability to track individual cell transcriptomes makes scRNAseq a powerful tool in understanding primordial follicle activation. Several studies have now used scRNAseq both in the cycling (21, 32) and ageing ovary (36, 37), where the focus is commonly on the oocyte. The transcriptomic datasets we provide here highlight the relevance of somatic cells in follicle activation. We also reveal *Slc18a2* as a novel primordial follicle activation gene. *Slc18a2* encodes Vesicular Monoamine Transporter 2 (VMAT2), a protein primarily characterised to transport monoamines into intracellular vesicles in the brain (38). VMAT2 is responsive to ovarian steroids (38) and expressed within rat granulosa cells (39) but is yet to have a clear function within primordial follicles. The upregulation of *Slc18a2* during primordial follicle activation may allow cuboidal granulosa cells to store monoamines, in preparation for gonadotropin-dependent folliculogenesis. Regardless, the identification of key changes in the transcriptome of pregranulosa cells provides further evidence that follicle activation is likely initiated within the pregranulosa cells (15). This is in agreement with studies illustrating activation changes in response to mechanical pressure (40) and metabolic changes (41). Consulting this dataset alongside previously published scRNAseq studies of the developing ovary will create a more comprehensive picture of granulosa cell changes across germ cell nest breakdown, primordial follicle formation and activation (5, 20, 21, 35, 42-44). Potential opportunities also exist to correlate this data with human ovary scRNAseq studies and those from other species to discover how mechanisms of follicle activation are conserved (22).

The bioinformatic approaches within the study provide new opportunities to discriminate transcriptionally similar cell types. Herein, we utilised two complementary approaches for dissecting activating pregranulosa cells from activated granulosa cells, namely Entropy Sorting (ES) and combining scRNAseq with bulk RNAseq from established infertility mouse models. As in previous uses of ES (30, 45, 46), our work demonstrates how the mathematical framework of ES can elucidate subtle changes in transcriptional profiles along developmental trajectories that conventional bioinformatics approaches may struggle to identify. Other developmental questions with dynamic and terminal trajectories could benefit from the application of this approach as a way of more accurately selecting gene targets underpinning development. The second bioinformatic approach was to functionally interrogate genes identified via ES using an infertile mouse model. We utilised a *Cdkn1b*^-/-^ mouse line that has an accelerated primordial follicle activation phenotype in the neonatal mouse ovary. Genes upregulated in *Cdkn1b*^-/-^ ovaries were part of the novel gene expression signature of activating pre-granulosa cells identified in our scRNAseq data, along with mature granulosa cell genes. The combination of scRNAseq data with bulk RNA-sequencing supports and strengthens our characterisation of the genes involved in primordial follicle activation, and as such we highlight this methodology as an under-utilised approach. This type of comparison has previously been informative in studies on embryonic somatic gonadal cells (47). In the context of this study, the combination of these bioinformatic tools was effective in identifying TNNI3 as a novel putative regulator of pregranulosa-driven primordial follicle activation.

*Tnni3* is an 8-exon gene located on chromosome 7 that encodes a 211-residue protein. TNNI3 functions in cardiac tissue as the inhibitory part of the three-subunit troponin complex and mutations in the gene in humans lead to cardiomyopathies (48). The function of TNNI3 in the ovary is currently unknown. A recent study of human growing follicles reported TNNI3 expression by granulosa cells and noted a characteristic ring-shaped staining pattern at the outer edge of these granulosa cells (31). Taken with our findings that *Tnni3* expression is upregulated in the granulosa cells of activating follicles, these data suggest a possible role of TNNI3 in inducing and maintaining the cuboidal shape of pregranulosa cells. Another explanation may be a requirement for TNNI3 to interact with protein kinase C (PKC), an established primordial follicle regulator, as TNNI3 is known to be phosphorylated by PKC in the mouse myocardium (49-51). Future work may provide a link between TNNI3 and other biological pathways, however conditional experiments will be required as *Tnni3*^-/-^ mice die perinatally from cardiac complications (52).

Questions remain as to whether the loss of cell-cycle inhibition in pregranulosa cells solely drives the precocious primordial follicle activation in *Cdkn1b*^-/-^ mutants. Evidence from a closer examination of *Cdkn1b*^-/-^ mutants suggests that germ cell nest breakdown and follicle formation may also be affected, as these mice form multi-oocyte follicles at higher rates than their control littermates (34). In this study, the phenotype of *Cdkn1b*^-/-^ mutants might also be exacerbated by the loss of p27^kip1^ in the oocytes, where expression comes on postnatally in wildtype ovaries. A conditional approach deleting *Cdkn1b* in pregranulosa cells specifically would provide a clear answer if primordial follicle activation is initiated in the pregranulosa cells and dysregulated by the absence of p27^kip1^. Regardless, it is clear from this body of work that further interrogation is necessary to find the master regulators of follicle activation.

In conclusion, this study presents transcriptomic datasets to interrogate primordial follicle activation in the mouse. We identify a gene expression signature within pregranulosa cells that are undergoing primordial follicle activation and highlight p27^kip1^-dependent genes within this signature. We utilise a recently described mathematical framework, Entropy Sorting, to discriminate activation-specific pregranulosa genes as high confidence imputed regulators of primordial follicle activation. Using this method, combined with an established infertile mouse model, we reveal a likely role for *Tnni3* as a regulator of activation within pregranulosa cells. Our data provide further evidence that it is the pregranulosa cell changes and not the oocytes, that underpin primordial follicle activation. Moreover, we hypothesise that the genes identified in Gc_3 by the Entropy Sort Score (ESS) are a critical part of the molecular control initiating this process. Our datasets, along with the bioinformatic approaches we have employed, will be valuable resources for the interrogation of mechanisms underlying mammalian follicle activation and female fertility.

## Materials and Methods

### Mice

All experiments carried out on mice were approved under the UK Animal (scientific procedures) Act 1986 (Project license 70/8560 and PP8826065). C57BL/6J females (RRID: IMSR_JAX:000664) were supplied by the Francis Crick Institute Biological Research Facility. *Cdkn1b*^*tm1Mlf*^ (*Cdkn1b*^-/-^) (RRID: IMSR_JAX:002781) (53) were maintained on a C57BL/6J/CBA mixed background and mated heterozygously. The Cdkn1b^tm1Mlf^ Foxl2^tm1(EGFP)Rlb^ (Cdkn1b^-/-^; Foxl2^P2A-eGFP/+^) were likewise maintained on a C57BL/6J/CBA mixed background and mated homozygously for the Foxl2^P2A-eGFP^ allele and heterozygously for the Cdkn1b^-/-^ allele (54).

### Tissue dissociation and scRNAseq library preparation

Ovarian tissue was dissociated as previously described (24). After mechanical dissociation, and filtration through a 20 µm pre-separation filter, approximately 10,000 live cells were used to generate libraries using the Chromium Single Cell 3’ kit v3.1, according to the manufacturer’s instructions (55). The libraries were quantified using the TapeStation (Agilent) to confirm quantity and purity, before sequencing on an Hiseq4000 (Illumina). Sequencing outputs were processed using the 10x CellRanger (version.3.0.2) to generate single-cell count data for each time point, using a mouse reference index provided by 10x Genomics (refdata-cellranger-mm10v3.0).

### Tissue dissociation and bulkRNA sequencing library preparation

Ovaries from *Cdkn1b*^+/+^ and *Cdkn1b*^-/-^ mice were collected at E13.5, E18.5 and PD8. A single ovary was dissociated as described above and RNA was extracted using the RNeasy Plus Micro kit (Qiagen). RNA quality was assessed the TapeStation (Agilent). Libraries were constructed using the NEBNext Low Input RNA Library Prep Kit (E6420L, New England Biolabs) according to the manufacturer’s instructions. All libraries were quantified using the TapeStation (Agilent) and sequenced on an Hiseq400 (Illumina) to achieve an average of 25 million reads per sample.

### Single-cell RNA sequencing analysis

Raw single-cell sequencing data from all three timepoints were processed using the 10x CellRanger pipeline (10x Genomics) and analysed with the Seurat package in R (56). All cells with fewer than 1500 RNA features were removed. Principal component analysis was performed on variable genes and 20 principal components were used to generate the UMAP plots using the default method from the Seurat package. Feature selection for UMAP generation was done using default seurat highly variable selection approaches (57) and continuous Entropy Sort Feature Weighting as previously described (30). The cESFW workflow used to identify the set of 4376 genes used to produce a high resolution UMAP embedding of pregranulosa cell differentiation may be found at the following GitHub repository, https://github.com/aradley/Taylor_Pregranulosa_Cells.

### Bulk RNA sequencing analysis

Libraries were sequenced on an Illumina HiSeq 4000 machine. The ‘Trim Galore!’ utility version 0.4.2 was used to remove sequencing adaptors and to trim individual reads with the q-parameter set to 20 (https://www.bioinformatics.babraham.ac.uk/projects/trim_galore/). The sequencing reads were then aligned to the mouse genome and transcriptome (Ensembl GRCm38release-89) using RSEM version 1.3.0 (58) in conjunction with the STAR aligner version 2.5.2 (59). The sequencing quality of individual samples was assessed using FASTQC version 0.11.5 (https://www.bioinformatics.babraham.ac.uk/projects/fastqc/) and RNA-SeQC version 1.1.8 (60). Differential gene expression was determined using the R-bioconductor package DESeq2 version 1.14.1 (61, 62). Log fold changes above 1 and below - 1 with a padj of less than 0.05 were considered significant.

### Data availability

The data supporting the findings of this study are available in the main text and the supplementary materials. Datasets have been submitted to NCBI GEO (GSE246098).

### Histology staining

Ovaries for hematoxylin and eosin (H&E) staining were fixed in chilled 4% (w/v) paraformaldehyde at 4 °C for 1 to 2 hrs. Following fixation, ovaries were washed in PBS 3 times for 1 hr each, before cryoprotection in 30% (w/v) sucrose in PBS overnight. The ovaries were then embedded in OCT mounting medium, frozen on an ethanol/dry ice slurry and stored at -80 °C. Samples were cryosectioned at 10 μm and sections were placed on Superfrost Plus glass slides and air dried for 5 minutes. H&E staining was performed by the Francis Crick Institute Experimental Histopathology facility. H&E images were taken at 10x magnification and 40x magnification, with scale bars of 100 µm and 25 µm respectively.

### Cryosectioning ovaries

Ovaries were collected from Foxl2^P2A-eGFP^ mice (54), fixed in 4% paraformaldehyde solution on ice for 30 mins followed by a further 1hr incubation at room temperature. Samples were subsequently cryopreserved 30% sucrose in PBS overnight at 4°C and embedded in OCT media prior to cryosectioning. OCT-embedded ovaries were serially sectioned into 12µm sections.

### Immunofluorescent staining of cryosectioned ovaries

Cryosectioned ovaries were washed in PBS, blocked with 10% donkey serum (blocking buffer) for 2hrs at room temperature. The primary antibody against TNNI3 (ab47003), FOXL2 (AB5096), and DDX4 (AB13840) were incubated at 4°C overnight at 1:200 dilution blocking buffer. Subsequent to washes, the secondary antibody (Donkey anti-Rabbit IgG Alexa Fluor 488 and Donkey anti-Goat Alexa Fluor 647) was diluted 1:500 in blocking buffer and incubated at room temperature for 2hrs. The nanobody against GFP (GBA-647n) was at 1:200 in blocking buffer for both primary and secondary incubations. Sections were then counterstained for 30 seconds with 4⍰,6-diamidino-2-phenylindole (DAPI) diluted 1:10000. Sections were mounted with Aqua-Poly/Mount and covered with glass coverslips. Imaging was done using a Zeiss Invert880 with Airyscan 2019 confocal microscope.

### Hybridization chain reaction (HCR)

HCR was conducted with reagents purchased from Molecular Instruments, Inc. Frozen ovarian sections were removed from -80°C storage and immersed for 5 minutes in 50%, 70% and 100% EtOH/H2O followed by immersion in PBS. A barrier was drawn around the tissue using a hydrophobic pen. Tissue was pre-hybridised by placing 100-200 μL of preheated (37°C) probe hybridization buffer directly on top of tissue and incubating for 10 minutes in a 37°C humidified chamber. HCR probe solution was made to the desired concentration by combining HCR probe stock with 37°C hybridisation buffer. The optimal probe concentrations were 128nM *Tnni3* probe (molecular instruments lot RTH757) and 4nM *Slc18a2* probe (molecular instruments lot RTH755). 25-30uL of probe solution was placed directly on top of each section and incubated for 16hrs at 37°C in a humidified chamber. Probe solution was removed by sequential 15 minute washes in 100%, 75%, 50% and 25% probe wash buffer diluted in a solution of 0.75M sodium-chloride 0.075M sodium-citrate with 0.1% Tween 20 (SSCT) at 37°C. Slides were then washed a further 15 minutes in 100% SSCT. Samples were then pre-amplified by adding 100-200uL of amplification buffer per slide and allowed to incubate for 30 minutes at room temperature. Hairpin solutions were made by snap cooling hairpin H1 and H2 at 95 ⍰C for 90 seconds and then cooling to room temperature in a dark drawer for 30 min. Hairpins were heated and snap cooled using a Biorad C1000 Touch Thermo Cycler. Snap cooled hairpins were then added to the amplification buffer to achieve a final concentration of 60nM. 25-30uL of hairpin solution was placed directly on top of each section and incubated for 16hrs at room temperature in a humidified chamber. Excess hairpins were removed via 2 x 30 min washes in SSCT. Sections were then counterstained for 30 seconds with DAPI diluted 1:10000. Sections were mounted with Aqua-Poly/Mount and covered with glass coverslips. Imaging was achieved using a Zeiss Invert880 with Airyscan 2019 confocal microscope.

## Acknowledgements and Funding Sources

## Acknowledgements

This work was supported by the Francis Crick Institute, which receives its core funding from Cancer Research UK (CC2116), the UK Medical Research Council (CC2116), and the Wellcome Trust (CC2116). We acknowledge the generous financial assistance to R.L.B. by the UK Biotechnology and Biological Sciences Research Council (BB/ N018680/1). We also gratefully acknowledge the financial assistance to J.M.S. and M.A.B. by the Australian National Health and Medical Research Council (G1600095) and the Hunter Medical Research Institute (G1501433 and G1801335). E.R.F. and B.N.C. are recipients of Australian Government Research Training Program Scholarships. We thank the Advanced Sequencing Facility at the Francis Crick Institute for their help in performing the sequencing studies in this manuscript. We would like to thank members of the Lovell-Badge and Sutherland laboratories for their comments on the manuscript.

## Author contributions

G.T. and E.R.F. conceived the study and wrote the manuscript. J.M.S., C.G., M.A.B. and R.L.B. provided expertise and feedback on the study design. All experiments were performed by E.R.F., G.T., B.N.C. and C.G.. RNAseq and scRNAseq analyses were performed by A.R. and S.B.. All the authors critically revised the manuscript. R.L.B. and J.M.S. provided resources for this work.

