## Supplemental Figures for "Pregranulosa cells engage a distinct transcriptional programme prior to cell-cycle dependent primordial follicle activation"

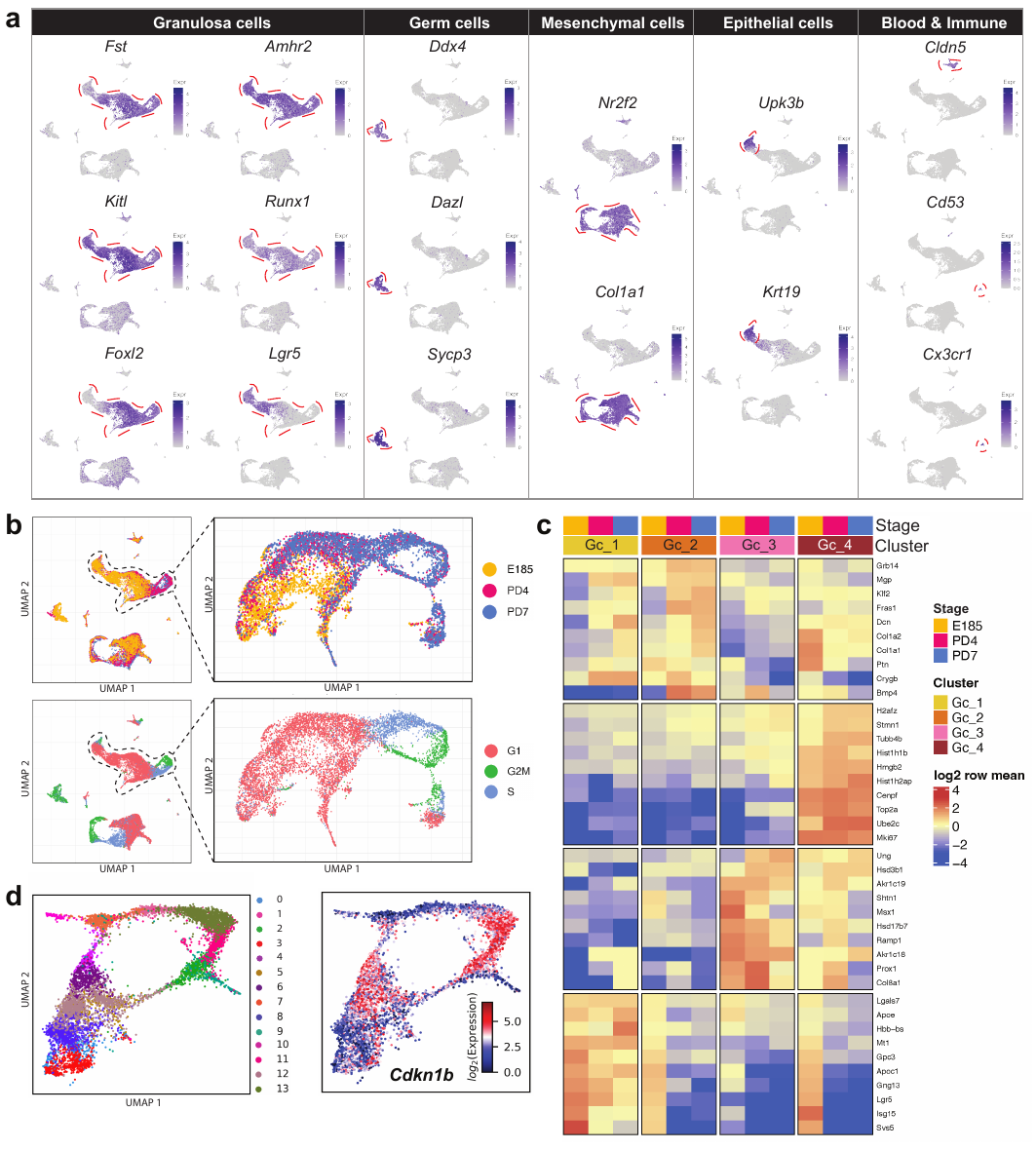

**Supplemental Figure 1. Identification of activating pregranulosa cell gene expression**

**signature in mouse ovaries.**

**a** UMAPs showing cell type markers for granulosa cells (*Fst*, *Kitl*, *Foxl2*, *Amhr2*, *Runx1*, *Lgr5*), germ cells (*Ddx4*, *Dazl*, *Sycp3*), mesenchymal cells (*Nr2f2*, *Col1a1*), epithelial cells (*Upk3b*, *Krt19*) and blood and immune cells (*Cldn5*, *Cd53*, *Cx3cr1*).

**b** UMAP of subclustered cells from Ep_1, Gc_1, Gc_2, Gc_3, Gc_4 and Gc_5 with three timepoints highlighted E18.5 (yellow), PD4 (magenta) and PD7 (blue). UMAP of subclustered cells from Ep_1, Gc_1, Gc_2, Gc_3, Gc_4 and Gc_5 with cell cycle phase of cells indicated: G1 (red), G2M (green) and S-phase (blue).

**c** Heatmap of top ten markers of Gc_1, Gc_2, Gc_3 and Gc_4 using Highly Variable Gene Selection.
**d** UMAP of Kmeans unsupervised clustering of scRNA-seq samples from Ep_1, Gc_1, Gc_2, Gc_3, Gc_4 and Gc_5 populations subsetted to the 4376 identified by cESFW reveals 14 distinct clusters. UMAP showing expression of dormancy marker *Cdkn1b*.

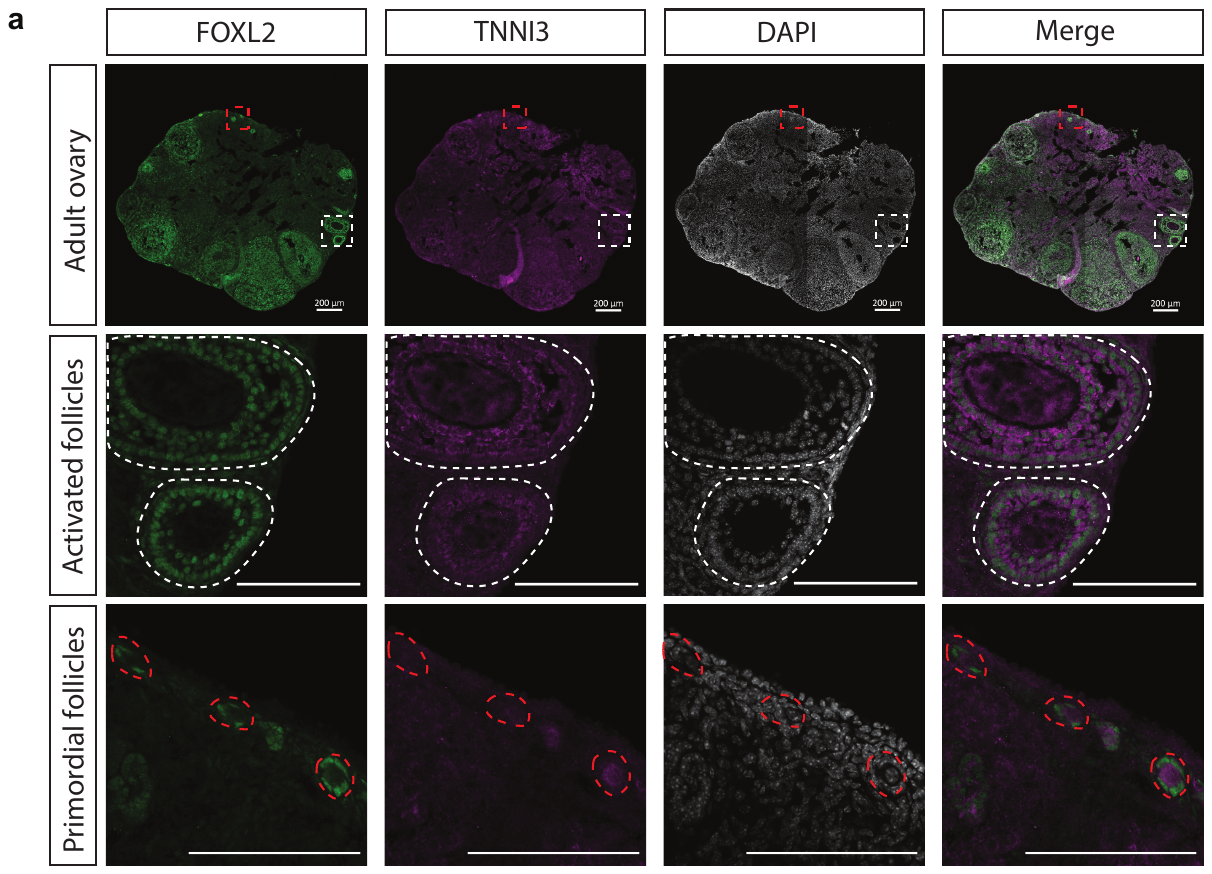

**Supplemental Figure 2. TNNI3 expression within granulosa cells of activated follicles in adult mouse ovaries.**

**a** Immunofluorescence on WT ovarian cryosections of 4 month old adult ovaries (n=3). FOXL2 (green) labelling all granulosa and pregranulosa cells, TNNI3 (purple) and DAPI (white). 10x magnification images are shown in the top row with dashed red and white coloured boxes highlighting two regions-of-interest (ROI) (scale bar 200µm). The white dashed ROI contains cortical activated follicles shown at 40x magnification (activated follicles). The red dashed ROI contains primordial follicles shown at 40x magnification (primordial follicles).

**Supplemental Figure 3. Disruption of p27^Kip1^ highlights functional aspects of activating pregranulosa cell gene expression signature.**

**a** Immunofluorescence staining on E18.5 from WT (n=3) and *Cdkn1b^-/-^* (n=3) ovaries. DDX4 staining of oocytes and FOXL2 staining of pregranulosa cells. Images at 40x (scale bars 50 µm) and 63x with 3x zoom (scale bars 10µm).

**b** PCA plots showing correlation of biological replicates from bulk RNA sequencing on WT and *Cdkn1b^-/-^* ovaries at E13.5, E18.5 and PD8.

**c** MA plot showing differential gene expression between WT and *Cdkn1b^-/-^* ovaries (upregulated genes in red, downregulated in blue, n = 4 replicates per group) at E18.5 and

PD8. Black dots indicate genes below the p ≤ 0.05 cutoff.

**Supplemental Dataset 1**

Differential gene expression analysis of cluster Gc_1 vs all other Gc clusters, presented by highest logFC.

**Supplemental Dataset 2**

Differential gene expression analysis of cluster Gc_2 vs all other Gc clusters, presented by highest logFC.

**Supplemental Dataset 3**

Differential gene expression analysis of cluster Gc_3 vs all other Gc clusters, presented by highest logFC.

**Supplemental Dataset 4**

Differential gene expression analysis of cluster Gc_4 vs all other Gc clusters, presented by highest logFC.

**Supplemental Dataset 5**

Entropy sorting rankings of all Gc clusters and Epithelial cluster 1 (Ep_1).

**Supplemental Dataset 6**

Differential gene expression analysis on *Cdkn1b^-/-^* ovaries vs WT ovaries at E13.5, E18.5 and PD8 timepoints.
